# Conditioning nausea in a laboratory setting: A pilot study

**DOI:** 10.1101/243121

**Authors:** Kathryne Van Hedger, Joel S. Cavallo, Nicholas A. Ruiz, Harriet de Wit

## Abstract

Cancer patients can experience nausea as they approach the place where they have received chemotherapy treatment. This nausea is likely the result of Pavlovian conditioning, where the previously neutral environment acquires conditioned properties, in this case conditioned nausea, because its association with feeling ill. To investigate this phenomenon under controlled conditions, we studied the acquisition of conditioned nausea using a distinct environment paired with an emetic drug in healthy young adults. We measured two indices of conditioning: i) conditioned place aversion, and ii) conditioned drug-like (nausea) responses. Healthy volunteers (N=32) first rated their preference for two testing rooms, and then underwent four conditioning sessions in which they received either syrup of ipecac (5 ml) or placebo. A Paired Group (PG; N=17) always received ipecac in their initially preferred room and placebo in the other, while an Unpaired Group (UG; N=15) received ipecac and placebo in both rooms. Conditioned responses were assessed with i) time spent in each room, ii) room liking and preference, and iii) feelings of nausea in each room. There was no evidence of conditioned place aversion as measured by either time spent or ratings of room liking. However, the PG did report a small increase in nausea in the ipecac-paired room. Although the conditioned responses in this study were not robust, this procedure is a first step towards studying conditioned aversive drug responses in humans, which will enable development of future studies to prevent or treat anticipatory nausea.

## Introduction

Cancer patients report anticipatory nausea when they approach locations where they have received chemotherapy treatment. This phenomenon is likely the result of Pavlovian conditioning, in which a previously neutral environment acquires conditioned properties (e.g., conditioned nausea) because of its association with feeling ill from the chemotherapeutic agent. Pavlovian conditioning has been studied extensively in laboratory animals and more recently in humans using the conditioned place preference procedure (Bardo & Bevins, 2000; Childs & de Wit, 2009, 2013; Philpot, Badanich, & Kirstein, 2003; Tzschentke, 2007). Most of these studies investigate drugs with rewarding properties, but other studies have demonstrated conditioned place aversion with drugs that have unpleasant effects including nausea (Parker & Mcdonald, 2000; Morrow, Arseneau, Asbury, Bennett, & Boros, 1982; Nesse, Carli, Curtis, & Kleinman, 1980; Redd & Andresen, 1981). That is, animals avoid an environment previously associated with emetic agents (Parker & Mcdonald, 2000). In addition to avoiding the place, these animals also exhibit specific conditioned drug-like effects such as ‘gaping’ when placed in the environment paired with an emetic drug (Parker, Rock, Sticht, Wills, & Limebeer, 2015; Rock, Limebeer, & Parker, 2015). Gaping is considered to be a sign of nausea in rats. In the present study, we designed a procedure to study the acquisition of conditioned place aversion with an emetic drug in human participants, using methods similar to those used in laboratory animals (e.g., Rock et al., 2015). We measured conditioning by assessing both aversion for the room where the drug was administered and subjective nausea ratings after conditioning.

Our goal was to develop a laboratory model for associative conditioning with an emetic drug in humans. Healthy volunteers received syrup of ipecac (IP; 5 ml) in one room and placebo (PBO) in another room, twice each on separate days. A paired group (PG), received IP consistently in one room, and an unpaired group (UG) received IP and PBO in both rooms. On a subsequent day (without drug administration) we assessed the amount of time subjects spent in each room, their liking of the two rooms, and feelings of nausea in both rooms. We hypothesized that participants in the PG would avoid, dislike, and exhibit nausea in the IP-paired environment.

## Materials and Methods

### Participants

Healthy volunteers (N=32) aged 18-35 years were recruited and screened with an in-person psychiatric interview, physical examination, electrocardiogram, and drug use history. Exclusion criteria included Major Axis I psychiatric disorders *(Diagnostic and Statistical Manual of Mental Disorders (5th ed.)*, 2013), serious medical conditions, current or past year substance dependence, and consumption of more than four alcoholic or caffeinated beverages/day. Women who were pregnant, planning to become pregnant or lactating were excluded. Participants had a minimum of a high school education, fluency in English, and BMI between 19 and 26 kg/m^2^. Participants provided written consent and were paid for their participation. This study was approved by the University of Chicago Biological Sciences Division Institutional Review Board.

### Study Design

This study used a six-session mixed within- and between-subjects design with two groups (PG, n=17; and UG, n=15). The first session provided baseline ratings of room preference. Then, during four conditioning sessions, subjects ingested IP or PBO in one of two distinctive rooms. The PG received IP twice in one room and PBO twice in the other room, whereas the UG received IP and PBO once each in both rooms. PG subjects were assigned to receive IP in their initially preferred room. The order of IP and PBO alternated between sessions and was counterbalanced between participants. On the last (post-conditioning) session participants rated and spent time in the two rooms. The primary outcome measures were: time spent in each room, room liking, and ratings of feeling nauseous.

### Session Procedures

*Session Locations:* Sessions were conducted in comfortably furnished rooms with movies and reading materials. Orientation and screening were conducted in a neutral room, and two rooms, designated Room A and Room B were used for conditioning. Rooms A and B were of similar size and lighting, and adjacent to each other, joined by a common hallway. The rooms differed in décor (color of the furniture, posters on the walls), and had distinctive scents (floral versus clean linen air freshener).

*Orientation/Pre-test Session:* During this session participants provided informed consent, were given instructions about drug abstinence before the sessions, and completed a pre-conditioning preference test. They were told that the study examined the relationship between drug effects and environment, and were informed that they could receive a drug that induces short-term nausea or vomiting. Participants explored the two testing rooms for 1 minute and their movements were videotaped. Afterwards, subjects rated their liking for each room and their preference between the two rooms. For the PG, these room preference ratings were used to determine which room would be paired with IP.

*Conditioning Sessions:* Each participant completed four 2-hour sessions from 9am to 11am, spaced 2-7 days apart. Upon arrival at every session, subjects were tested for compliance with drug use restrictions using breathalyzers (measuring blood-alcohol levels; Alco-sensorIII, Intoximeters, St Louis, MO), urine drug tests (ToxCup, Branan Medical Corporation, Irvine CA), and women were tested for pregnancy (AimStickPBD, hCG professional, Craig Medical Distribution, Vista, CA). Then they were escorted to the designated room (A or B) where baseline blood pressure (BP) and heart rate (HR) measures were obtained and subjects completed a baseline “brief nausea questionnaire” (BNQ) that asked about feelings of nausea, queasiness, and urge to vomit at that moment. At 9:20am, IP or PBO was administered under double-blind conditions with 100 ml of water. IP (5 ml; Lifestyle Pharmacy, Windsor, ON) was mixed with 5 ml of Ora-Sweet syrup (Paddock Laboratories, Minneapolis, MN). PBO syrup consisted of 10 ml of Ora-Sweet alone. BNQ measures were assessed at eight times throughout the 2 hour session. Physiological measures were obtained at baseline and 20, 50, and 110 minutes after drug administration. At the end of the session, participants rated the strength of the nausea experienced during the session and the number of times vomited.

*Post-test Session:* This session was conducted 2-7 days after the last conditioning session. During the post-test, participants completed the same measures as during the pre-test. They explored both rooms for 1 minute and then spent 30 seconds in each room completing ratings of liking and subjective effects.

### Drug Effect Measures

*Cardiovascular and Subjective Effects during Conditioning Sessions:* HR and BP were measured periodically during the conditioning sessions. Mean arterial pressure (MAP) was calculated using the formula: [systolic BP+2 x diastolic BP]/3. Subjective nausea effects were measured throughout each conditioning session using the BNQ.

### Dependent Measures

*Room Exploration Time:* Participants were videotaped for 1 minute while exploring the testing rooms during the sessions before and after conditioning. Blinded research assistants recorded the total time spent in each room.

*Self-reported “Liking” and Room Preference:* During the pre-test session and the post-test session, subjects rated their liking of the rooms and which room they preferred using 100mm visual analog scales. For liking, they marked a vertical line indicating how much they liked each room (A and B) on two scales anchored with ‘Dislike Very Much’ and ‘Like Very Much.’ For preference, they marked their relative liking of the rooms on a 100 mm line anchored with ‘Prefer Room A’ and ‘Prefer Room B.’

*Brief Nausea Questionnaire (BNQ):* The BNQ consisted of 100 mm visual analog scales assessing feelings of nausea at the moment from ‘Not at all’ to ‘Very Much.’ Adjectives included feelings of nausea, headache, dizziness, queasiness, urge to vomit, tiredness, anxiety, sweatiness, and salivation. During the post-test session, subjects completed the BNQ once in each room after sitting in each room for 30 seconds.

### Data Analysis

*Data Processing:* Data were analyzed using SPSS statistical software (version 22, SPSS Inc., IBM, Chicago, IL). Before conducting analyses, we confirmed that overall participants did not prefer one room (A or B) over the other before conditioning. Values for each time point and area under the curve (AUC) values were averaged within IP sessions and within PBO sessions for all participants. The PG and UG groups did not differ in demographic characteristics.

*Drug Effects Measures:* BNQ scores and cardiovascular measures (HR, MAP) were analyzed using AUC scores during conditioning sessions. AUC values were calculated using the trapezoid method relative to the baseline. These values were then analyzed using independent samples *t*-tests to examine differences between the PG and UG.

*Dependent Measures*: Room exploration was analyzed using a repeated measures ANOVA of change scores for exploration (post-test minus pre-test) for each room (initially preferred, initially non-preferred) as a within-subjects factor and group (PG, UG) as a between subjects factor. Room liking was analyzed similarly to room exploration using repeated measures ANOVA. Change in room preference from before to after conditioning was analyzed using a Pearson chi-squared to compare changes in preference in the PG and UG. Context-produced subjective effects at the post-test session were analyzed using paired t-test to compare subjective effect scores for nausea, queasiness, and urge to vomit between the preferred and non-preferred rooms for each group.

## Results

### Demographic Characteristics

The PG and UG did not differ in age, sex, BMI, or years of education completed. Most participants were in their mid-20s and Caucasian (Table 1).

**Table 1:** Demographic Information and Lifetime Nonmedical Drug Use (N=32).

| <u>Category</u> | <u>Paired Group</u> | <u>Unpaired Group</u> |
| --- | --- | --- |
| *Gender* |  |  |
| Male/Female | 8/9 | 7/8 |
| *Age* | 24.82 (0.19) | 24.33 (0.21) |
| *Education (years)* | 15.44 (0.15) | 15.07 (0.06) |
| *BMI* | 22.03 (0.09) | 22.61 (0.12) |
| *Race* |  |  |
| Caucasian | 64.7% (11) | 66.7% (10) |
| African-American | N/A | 26.7% (4) |
| Asian | 17.6% (3) | N/A |
| Other | 11.8% (2) | 6.7% (1) |
| Unknown | 5.9% (1) | N/A |
| *Lifetime Non-Medical Drug Use* |  |  |
| Marijuana | 76.5% (13) | 80.0% (12) |
| Opiates | 17.6% (3) | 33.3% (5) |
| Stimulants | 41.2% (7) | 53.3% (8) |
| Hallucinogens | 29.4% (5) | 60.0% (9) |
| MDMA | 35.3% (6) | 46.7% (7) |
| Sedatives | 17.6% (3) | 26.7% (4) |

The groups did not differ significantly on any measure. BMI = body mass index. Values indicate N (gender), Mean and SEM (age, number of years of education, BMI), or percent of participants and N (race and nonmedical drug use). N/A indicates no participants satisfied this category.

### Effect of IP

During the conditioning sessions, IP increased nausea, MAP and HR in both groups. Participants reported significant increases in nausea after IP compared to PBO (t(31) = 9.77, *p* < 0.001; Figure 1) and 29 out of 32 subjects vomited. IP significantly increased MAP and HR compared to PBO (t(31) = 4.67, *p* < 0.001 and *t*(31) = 5.37, *p* < 0.001; Figure 2). Subjective, emetic and cardiovascular effects during conditioning sessions were unrelated to body weight or sex.

**Figure 1:**
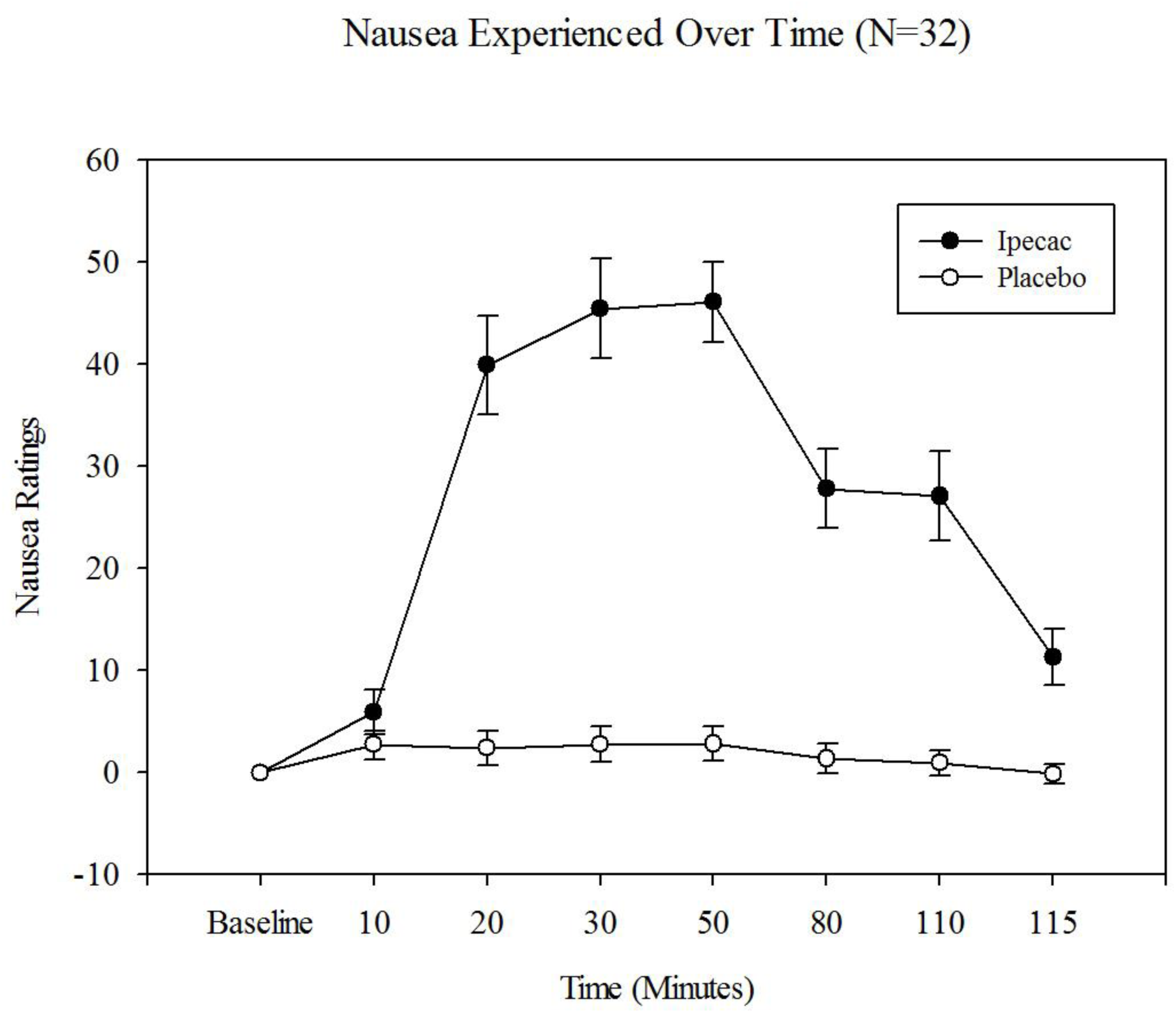
Mean nausea ratings (SEM) at each time point during the sessions, during the ipecac (IP, black circles) and placebo (PBO, white circles) sessions. Data for all 32 subjects were combined, as there were no differences between the groups or between the two administrations of IP. Baseline refers to the rating before receiving a drug, and error bars are standard error values. IP increased ratings of nausea and PBO did not.

**Figure 2:**
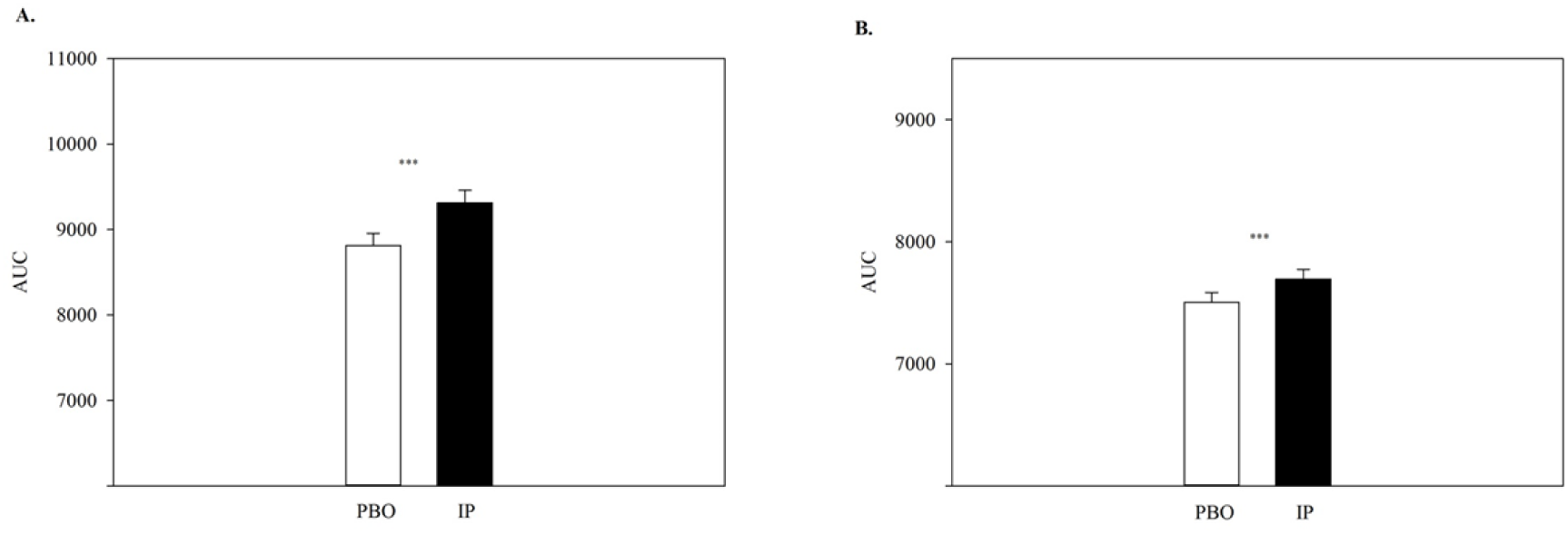
Mean area under the curve for mean arterial pressure (MAP, panel A) and heart rate (HR, panel B) during placebo (PBO) sessions (black bars) and ipecac (IP) sessions (gray bars) during conditioning sessions for all subjects taken together (N=32). Responses to IP did not differ across the two administrations, or between the PG and UG groups. IP significantly increased both MAP and HR compared to PBO.

### Dependent Measures

*Room Exploration Time:* There was no change in the amount of time subjects spent exploring the two rooms from pre- to post-test in either group. Thus there was no evidence of conditioned place aversion in the PG.

*Self-reported “Liking” and Room Preference:* Ratings of the initially preferred room declined after conditioning in both groups [significant main effect of room, F(1,30) = 7.78, *p* = 0.009 (Figure 3)], and this change was similar in the PG and UG.

**Figure 3:**
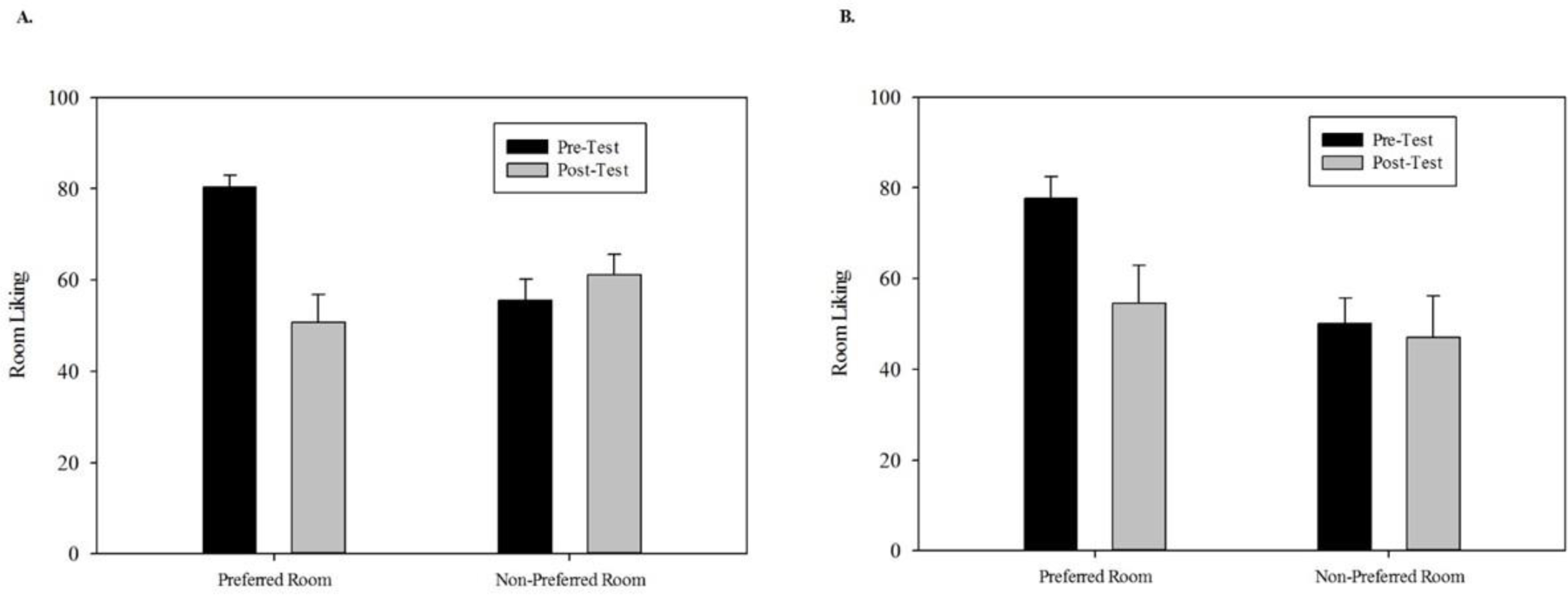
Mean room liking score for the initially Preferred and Unpreferred Room before (black bars) and after (grey bars) conditioning sessions in the Paired (N=17, panel A) and Unpaired (N=15, panel B) Groups. Participants rated liking of each room on a 100 mm line, labeled ‘not at all’ (0) to ‘extremely’ (100). Error bars are standard error values. After conditioning, liking scores of the initially preferred room decreased in both groups, but liking scores of the initially nonpreferred room remained stable.

*Room-Specific Subjective Effects at Post-Test:* At the post-conditioning session, the PG group reported more nausea in the IP-paired room (mean 2.62 ± 1.40 SEM) than the PBO-paired room (mean 1.04 ± 0.79 SEM; t(16) = 2.19, *p* = 0.04), but the effect was small (Cohen’s *d* = 0.25). Queasiness and urge to vomit were rated equally in the two rooms. The UG did not report differences in nausea, queasiness, or urge to vomit between the two rooms.

## Discussion

The present study is a first step toward developing a laboratory model for studying aversive drug conditioning in humans. Cancer patients report feeling nauseous as they approach the place where they received chemotherapy, and rodents exhibit conditioned place aversion and conditioned drug-like ‘gaping’ when they are placed in a context where they experienced an aversive event. In the present study IP reliably produced nausea and vomiting and increased heart rate and blood pressure. However, we found little evidence of conditioned place aversion. The PG did not avoid the room in which they had received IP, and their ratings of room liking or preference were not different from the UG. The PG reported a small, but significant, increase in conditioned nausea in the room paired with IP, a finding consistent with preclinical research of conditioned drug ‘gaping’ after an emetic drug (Parker et al., 2015; Rock et al., 2015).

There are several possible reasons why a conditioned place aversion did not develop and conditioned nausea was mild in the present study. It is probable that more drug pairings and more distinctive environments are needed to establish robust conditioned nausea in healthy individuals. It could also be the case that the physical features of the two rooms, located in adjacent rooms in a hospital corridor, were too similar to establish differential conditioning. In cancer patients the stimuli that comprise the conditioning context (i.e., a hospital with distinctive sights, sounds, smells, memories) provide a highly salient conditioned stimulus. This salience is likely compounded by the cognitive and emotional state of the patient receiving treatment (Morrow et al., 1982; Nesse et al., 1980; Redd & Andresen, 1981). In contrast, the contextual differences between our two conditioning rooms were minor. It is also possible that more pairings or stronger sickness experiences would lead to more substantial conditioning. Cancer patients experience numerous chemotherapy sessions with debilitating nausea, and there is evidence that anticipatory nausea in cancer patients is positively related to the number of chemotherapy sessions (Morrow et al., 1982; Nesse et al., 1980). Additionally, the chemotherapy sessions are separated by weeks, not days, and there is no placebo control condition in the patients.

Other aspects of our design and methods might have prevented robust conditioning. For example, the use of a biased design in which IP was paired with the participant’s initially preferred room may have interfered with the acquisition of conditioning. Some participants reported strongly preferring one room before conditioning, which might be difficult to reverse. Finally, participants’ knowledge that they would be receiving either IP or PBO at each session might have reduced the aversiveness of the IP-induced nausea or interfered with the acquisition of the conditioned association.

Despite these limitations, this study is a first step toward developing a laboratory paradigm to model aversive conditioning with emetic drugs in humans. Anticipatory nausea is a pervasive problem among patients undergoing chemotherapy, and it is refractory to current antiemetic treatments like ondansetron (Hickok et al., 2003; Morrow et al., 1997; Tyc, Mulhern, Barclay, Smith, & Bieberich, 1997). There is promising preclinical evidence suggesting that certain drugs preferentially target anticipatory nausea, including drugs that act on the endogenous cannabinoid system (Parker, Rock, Sticht, Wills, & Limebeer, 2015; Rock, Limebeer, & Parker, 2014). A human conditioned nausea paradigm will be critical for testing the effectiveness of these drugs, and will allow researchers to investigate possible pharmacological or behavioral treatments to reduce anticipatory nausea in cancer patients.

